# Metformin enhances anti-mycobacterial responses by educating immunometabolic circuits of CD8^+^ T cells

**DOI:** 10.1101/2020.08.26.269217

**Authors:** Julia Böhme, Nuria Martinez, Shamin Li, Andrea Lee, Mardiana Marzuki, Anteneh Mehari Tizazu, David Ackart, Jessica Haugen Frenkel, Alexandra Todd, Ekta Lachmandas, Josephine Lum, Foo Shihui, Tze Pin Ng, Bernett Lee, Anis Larbi, Mihai G Netea, Randall Basaraba, Reinout van Crevel, Evan Newell, Hardy Kornfeld, Amit Singhal

## Abstract

Diabetic patients taking metformin have lower risk for *Mycobacterium tuberculosis* (*Mtb*) infection, progression from infection to tuberculosis (TB) disease, TB morality and TB recurrence. However, a detailed mechanistic understanding of metformin’s protective immunological benefits on host resistance to TB is lacking. In this study, using mass cytometry we show that metformin treatment expands memory-like antigen-inexperienced CD8^+^CXCR3^+^ T cells in naïve mice, and in healthy and diabetic humans. Metformin-educated CD8^+^ T cells have increased (i) mitochondrial mass, oxidative phosphorylation, and fatty acid oxidation; (ii) survival capacity; and (iii) anti-mycobacterial properties. CD8^+^ T cells from *CXCR3^−/−^* mice did not exhibit metformin-mediated metabolic programming. In BCG-vaccinated mice and guinea pigs, metformin enhanced immunogenicity and protective efficacy against *Mtb* challenge. Collectively, our results demonstrate an important role of CD8^+^ T cells in metformin-derived host metabolic-fitness towards *Mtb* infection.

## Introduction

Tuberculosis (TB) remains a major threat to global health owing to the inadequacies of antimicrobial treatment, the lack of a highly effective preventive or therapeutic vaccine, and the rise of drug-resistant *Mycobacterium tuberculosis* (*Mtb*) infection. Recent interest has focused on a new therapeutic paradigm, host-directed therapies (HDT) for TB, to boost the antimicrobial efficacy of innate and adaptive immunity to TB while limiting excessive inflammation and tissue damage ^1, 2^.

Previously we discovered that type 2 diabetes mellitus (T2D) patients using metformin have reduced risk of severe TB disease ^3^ and *Mtb* infection ^4^. Metformin also enhanced the efficacy of anti-TB drugs, ameliorated lung pathology and reduced inflammation in *Mtb*-infected mice ^3^. Retrospective data from 8 clinical cohorts comprising a combined >250,000 individuals treated with metformin for T2D showed that metformin reduces the risk for progression to TB disease, mortality, lung cavitation and relapse, and that it accelerates sputum conversion independently of glycemic control ^3, 4, 5, 6, 7, 8, 9^. Metformin is a top candidate in the TB-HDT pipeline, but a detailed mechanistic understanding of its protective immunometabolic benefits is lacking.

While CD4^+^ T cells are generally recognized as the predominant effectors of adaptive immunity to TB, CD8^+^ T cells are also recognized to play a critical role in determining whether the *Mtb* infection is contained or progresses to active disease ^10^. Upon infection, naïve CD8^+^ T cells (T_N_) differentiate into antigen (Ag)-specific effector T cells (T_E_) and central memory T cells (T_CM_). The latter have the ability to survive, expand and generate cytotoxic CD8^+^ T_E_ cells upon encounter with a cognate Ag later in life ^11^. These T_CM_ are the major memory-like T cells in an infected or an immunized host. In contrast, in naïve mice housed under specific-pathogen free conditions, Ag-inexperienced memory-like CD8^+^ T cell (TM) population, sometimes known as virtual-memory (T_VM_) or innate-memory CD8^+^ T cells, have been described ^12, 13, 14^. These Ag-inexperienced CD8^+^ T_M_ cells (i) display unique and similar phenotypes to Ag-experienced CD8^+^ T_CM_ cells ^13, 14, 15, 16^, (ii) rapidly respond to primary antigenic stimuli ^14^, and (iii) mediate the protective immunity via non-canonical effector functions ^17^.

Survival, activation and effector function of T cells is fundamentally linked to cellular metabolic programming ^18^. T_E_ cells predominantly use glycolysis while T_CM_ cells use oxidative phosphorylation (OXPHOS) to meet energy demands ^18, 19^. CD8^+^ T_CM_ exhibit increased mitochondrial fatty-acid oxidation (FAO), spare respiratory capacity (SRC) and biogenesis ^20^. SRC is the reserve capacity to produce energy in the mitochondria beyond the basal state. The increase in FAO required for optimal CD8^+^ T_CM_ generation and survival depends on the mitochondrial import of long chain fatty acids ^20, 21^. Of note, CD8^+^ T_CM_ cells have been recently demonstrated to have lower SRC than CD8^+^ T_VM_ cells ^22^.

In this study, we sought to determine whether metformin’s mediated metabolic reprograming by metformin empowers CD8^+^ T cells to contain *Mtb* infection. We show that metformin treatment (i) expanded memory-like CD8^+^CXCR3^+^ T cells in naïve and in healthy and diabetic humans, (ii) induced mitochondrial SRC and FAO in CD8^+^ T cells, and (iii) enhanced BCG vaccine-elicited CD8^+^ T cell responses and efficacy in mouse and guinea pig TB models. Metformin-educated Ag-inexperienced CD8^+^ T_M_ cells showed gene expression signature of an activated T cell, and restricted *Mtb* growth in T cell deficient mice.

## Results

### Metformin-educated CD8^+^ T cells restrict M. tuberculosis replication

We previously reported that metformin attenuates immunopathology and reprograms CD4^+^ and CD8^+^ T cell responses in tissues of *Mtb*-infected wild-type C57BL/6 (WT) mice ^3^. To dissect the relative importance of metformin-educated CD4^+^ *vs* CD8^+^ T cells in TB defense, we compared their capacity to protect against *Mtb* infection in an adoptive transfer model. Splenic CD4^+^ or CD8^+^ T cells from WT mice treated with metformin or not (control group) were isolated and adoptively transferred into irradiated recipients, which were then infected with *Mtb* (Fig. 1a). Irradiated naïve recipients that did not receive any T cells were also infected (no transfer group). In two independent experiments we found that mice which received metformin-educated CD8^+^ T cells had 0.5 log10 reduced lung bacterial load at 21 days post-infection (p.i.) compared to the mice that did not receive any cells (Fig. 1b, c). At the same time, we noticed a 0.3 log10 reduction in the lungs of mice receiving metformin educated CD8^+^ T cells compared to those receiving CD8^+^ T cells from untreated mice (Fig. 1b, c). In these experiments, CD4^+^ T cells from metformin-treated and untreated donors failed to mediate the protection against *Mtb* (Fig. 1b, c).

**Figure 1.**
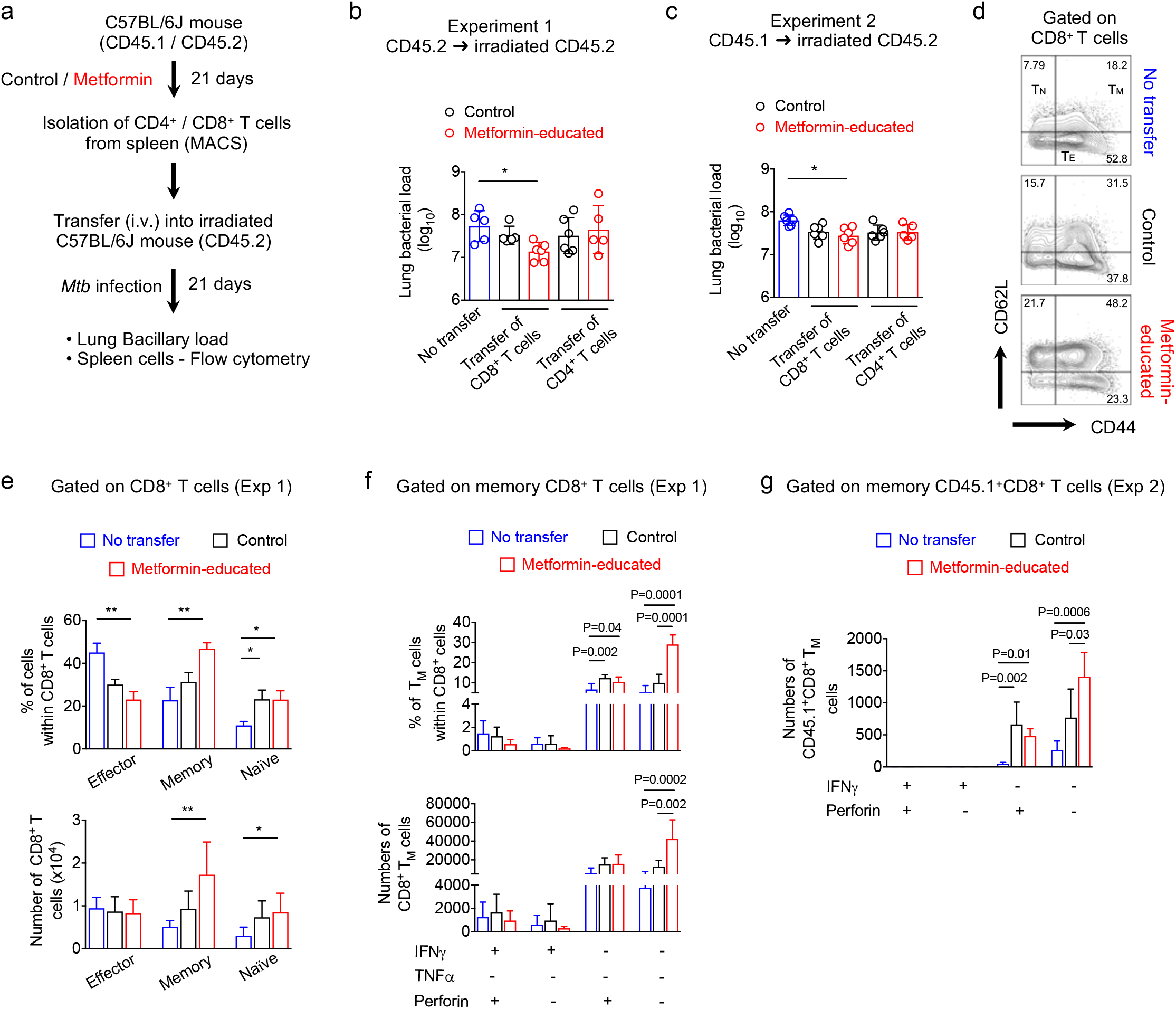
Metformin-educated CD8^+^ T cells protect against *M. tuberculosis*. **a** Experimental strategy and assessed readouts. **b, c** Bacterial load in the lung of recipient mice 21 days post-transfer, from two different experiments. No transfer – Irradiated recipients which did not receive any cells. Metformin-educated - CD4^+^ or CD8^+^ cells from donor mice treated with metformin; Control – Cells from untreated mice. **d** Flow cytometric analysis of splenic CD8^+^ T cells from mice of experiment 1 (b). T_M_: memory T cells, CD44^+^CD62L^+^; T_E_: effector T cells, CD44^+^CD62L^−^; T_N_: naïve T cells, CD44^−^CD62L^+^. **e** Frequency and number of CD8^+^ T_E_, T_M_, and T_N_ cells in the recipient mice of experiment 1. **f** Frequency and number of IFNg^+^, TNFa^+^ or Perforin^+^ CD8^+^ T_M_ cells from e. **g** Frequency and number of IFNg^+^ and/or Perforin^+^ CD45.1^+^CD8^+^ T_M_ cells in the recipient mice of experiment 2. Data represent mean ± SD of 5 - 6 mice per group. \**p* ≤ 0.05, \*\**p* ≤ 0.01 by One way ANOVA.

The reduction in the lung bacterial load in recipients of metformin-educated CD8^+^ T cells was paralleled by an increased frequency and total numbers of splenic memory CD8^+^CD44^+^CD62L^+^ T cells and naïve CD8^+^CD44^−^CD62L^+^ T cells, and decreased effector CD8^+^CD44^+^CD62L^−^ cells compared to the mice that did not receive CD8^+^ T cells (Fig. 1d, e and Supplementary Fig. 1a). Of note, there was no change in the total number of CD3^+^, CD4^+^ and CD8^+^ T cells in the recipient mice among different groups (Supplementary Fig. 1b). These results suggested that the predominant effect of metformin treatment in donor mice was on the memory compartment of CD8^+^ T cells over effector compartment, since more memory cells were observed within the total CD8^+^ splenic T cells from *Mtb*-infected recipients receiving metformin educated CD8^+^ T cells. In contrast, the proportions of memory to effector CD8^+^ T cells were equal in the spleens of *Mtb*-infected mice that received control CD8^+^ T cells (Fig. 1e). Mice receiving either metformin-educated or control CD8^+^ T cells had increased frequencies of perforin-expressing (Perforin^+^IFNγ^−^ TNFα^−^) memory CD8^+^ T cells compared to the mice that did not receive any T cells (Fig. 1f). In congenic recipients (Exp 2, receiving either metformin-educated or control CD8^+^ T cells) we saw similar increased numbers of Perforin^+^IFNγ^−^ donor memory CD45.1^+^CD8^+^ T cells compared to the mice that did not receive any T cells (Fig. 1g and Supplementary Fig. 1c). Surprisingly, mice receiving metformin-educated CD8^+^ T cells had increased numbers of double- or triple-negative memory CD8^+^ T cells (Fig. 1f, g), indicating perforin-, IFNγ- and TNFa-independent *Mtb* killing property of metformin-educated CD8^+^ T cells. CD4^+^ T cells have been reported to have such non-canonical anti-*Mtb* property ^23^. Overall, these results demonstrated the anti-*Mtb* activity of metformin-educated CD8^+^ T cells and suggested the potential of metformin to induce the expansion of memory CD8^+^ T cells with a unconventional effector phenotype.

### Metformin expands CD8^+^CXCR3^+^ T_M_ cells in naïve mouse tissues

To understand the effects of metformin exposure on CD8^+^ T cells in naïve mice, we investigated the phenotype of metformin-educated splenic CD8^+^ T cells by mass cytometry (CyTOF) ^24, 25^ using a panel of 40 markers (Fig. 2a and Supplementary Table 1). Analysis of live CD45^+^CD19^−^ TCRβ^+^CD90^+^CD4^−^CD8^+^ spleen cells (Supplementary Fig. 2a) using the nonlinear dimensionality reduction technique t-SNE with a machine-learning clustering algorithm (Phenograph) identified 20 distinct cell clusters with different expression characteristics of surface markers and intracellular transcription factors (Fig. 2b, c and Supplementary Fig. 2b). Based on the expression of CD62L and CD44, 20 clusters of CD8^+^ T cells were broadly classified into T_N_, T_E_ and T_M_ cells (Fig. 2b and Supplementary Fig. 2b). Automated machine-learning gating showed an expansion in the proportion of cells in clusters 9, 12 and 18, which comprised all TM cells, within the total metformin-educated CD8^+^ T cell population (Supplementary Fig. 2c). These clusters were strongly positive for conventional and unconventional memory T cell markers including CD62L, CD44, CD127, CD27, CD122, and Eomes (Fig. 2c and Supplementary Fig. 2b). They also expressed Tbet, CXCR3 and CD49d (Fig. 2c and Supplementary Fig. 2b), indicating an effector phenotype and differentiating them from CD8^+^ T_VM_ cells ^15, 26^. Tbet directly transactivates CXCR3 ^27^ and retroviral-mediated CXCR3 expression in *Tbet^−/−^* CD8^+^ T cells reconstitutes their ability to infiltrate inflamed tissues ^28^. CXCR3 plays an important role in T cell trafficking and function ^26^ and expression of CXCR3 on CD8^+^ TM cells is critical for populating the airways under steady state conditions ^29^.

**Figure 2.**
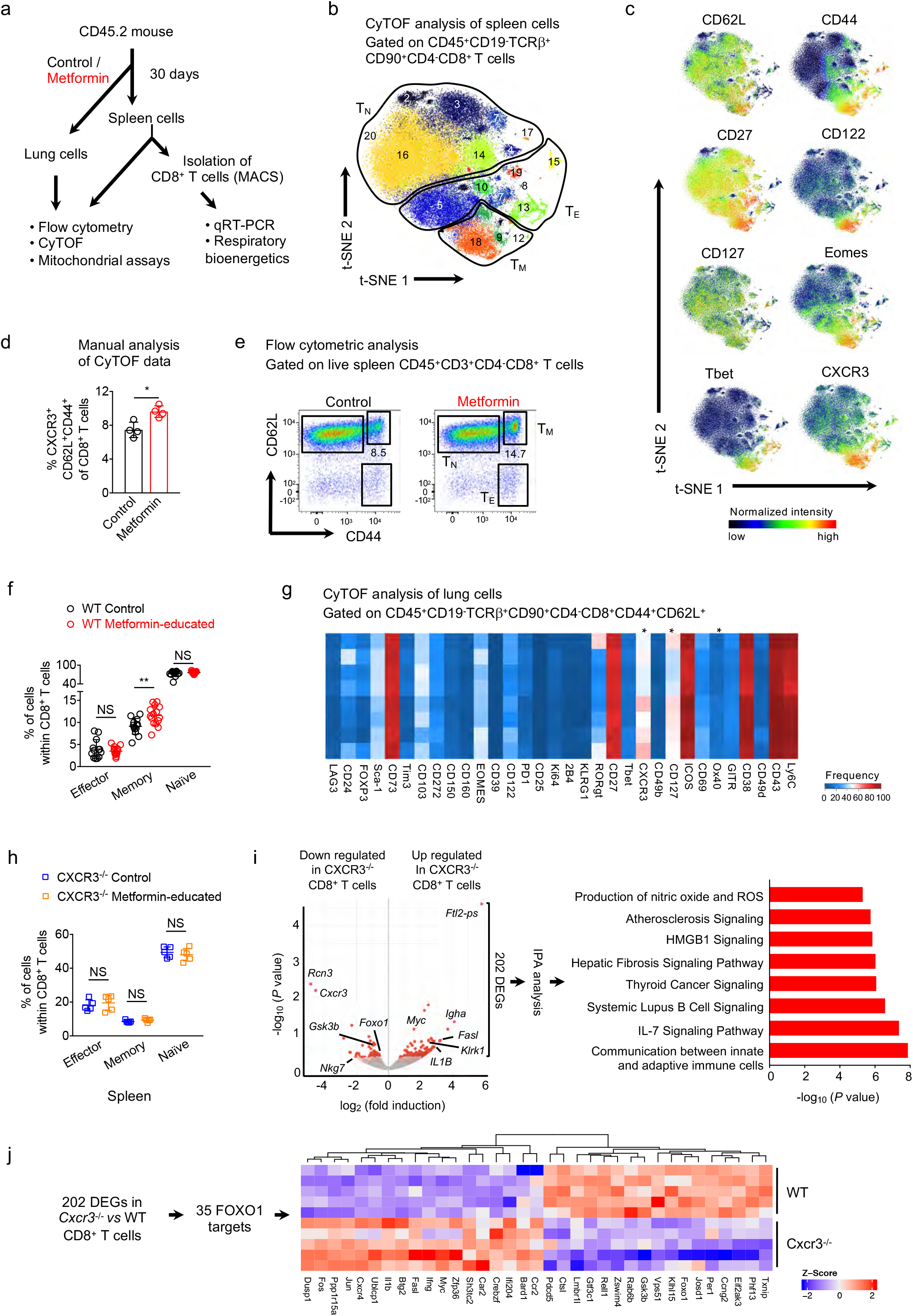
Metformin treatment expands CD8^+^CXCR3^+^ TM cells in spleen and lung of mice. **a** Experimental set-up of *in vivo* experiments. **b** Visualized t-SNE map of CD8^+^ T cells from spleen of wild type (WT) mice with automated cluster gating. t-SNE was performed on live CD45^+^CD90^+^TCRαβ^+^CD4^−^CD8^+^ cells after gating out B cells. **c** Normalized expression intensities of indicated marker was calculated and overlaid on the t-SNE plot, characterising all clusters of CD8^+^ T cells in spleen. **d** Manual gating analysis of CD8^+^CXCR3^+^ T_M_ cells in control and metformin-treated WT mice. **e, f** Flow cytometric analysis of splenic CD8^+^ T cell subsets from control and metformin-treated WT mice. Pooled data from three independent experiments. Means ± SD of 13-15 mice per group. **g** Heat map displaying frequency of various surface markers in lung CD8^+^ T cells of control and metformin-treated WT mice. N=4. \**p* ≤ 0.05 by two-tailed paired *t* test. **h** Flow cytometric analysis of spleen CD8^+^ T_E_, T_M_, and T_N_ cells of control and metformin-treated *CXCR3^−/−^* mice. Data shown one out of two experiments, mean ± SD of 5 mice per group. **i** Left - Volcano plot showing DEGs from RNA sequencing data of CD8^+^ T cells from *Cxcr3^−/−^* and WT mice. Right – Canonical pathway enrichment via Ingenuity pathway analysis (IPA) of 202 DEGs. **j** Heat map displaying the expression level of 35 FOXO1 targets in data shown in i. Data in d and f, \**p* ≤ 0.05, \*\**p* ≤ 0.01, Mann-Whitney *U* test. NS - Not significant.

Using a manual gating strategy, we validated the enrichment of CD8^+^ TM cells identified by machine-learning strategy, confirming an increased frequency of memory-like CD8^+^CXCR3^+^ cells after metformin treatment (Fig. 2d). The enhanced CD8^+^ T_M_ frequency induced by metformin was further confirmed by flow cytometry of splenocytes from naïve mice (Fig. 2e, f). Flow cytometric analysis also showed an enhanced frequency of total CD8^+^CXCR3^+^ cells in the spleen of metformin-treated naïve mice (Supplementary Fig. 2d). Interestingly, OVA protein-stimulated CD8^+^ OT-I cells cultured with IL-15 (a cytokine known to stimulate T_M_ cell differentiation ^30^) showed increased T_CM_/T_E_ ratio in the presence of metformin (Supplementary Fig. 3a, b), further demonstrating effectiveness of metformin in expanding the CD8^+^ T_CM_ compartment as well.

We next used CyTOF to decipher the heterogeneity in the lung CD8^+^ T cells from metformin-treated naïve WT mice (Supplementary Fig. 4a). Machine learning analysis showed metformin mediated increased expression of CXCR3 on lung (i) total CD8^+^ T cells (Supplementary Fig. 4b), (ii) CD8^+^ T_E_ cells (Supplementary Fig. 4c) and CD8^+^ T_M_ cells (Fig. 2g). This was further confirmed by manual gating of lung CyTOF data (Supplementary Fig. 4d) and flow cytometric analysis of lung cells (Supplementary Fig. 4e) from metformin-treated naïve mice. To test if CXCR3 was involved in the metformin-mediated expansion of CD8^+^ T_M_ cells in the spleen and lung, we compared untreated and metformin-treated *Cxcr3^−/−^* mice. The *Cxcr3^−/−^* mice had increased CD8^+^ T_E_ and reduced CD8^+^ T_M_ cells compared to WT mice in spleen (Fig. 2h) and lung (Supplementary Fig. 4f), confirming an earlier report ^31^, while in the present study metformin treatment did not alter the percentage of T_E_, T_M_ or T_N_ cells in *Cxcr3^−/−^* mice.

To determine why metformin failed to expand CD8^+^ T_M_ cells in *Cxcr3^−/−^* mice, we performed RNA sequencing on splenic CD8^+^ T cells. We found 202 differentially expressed genes (DEGs) in CD8^+^ T cells from *Cxcr3^−/−^* compared to WT mice (Fig. 2i, left panel and Supplementary Table 2). Ingenuity pathway analysis (IPA) of these DEGs revealed an enrichment of genes associated with communication between innate and adaptive immune cells and IL-7 signaling pathway (Fig. 2i, right panel and Supplementary Table 3). For optimal CD8^+^ T_M_ cell homeostasis a discontinuous IL-7 signaling has been proposed ^32^. Among DEGs, we found downregulation of *Foxo1* (Forkhead Box O1), *Gsk3b* (Glycogen synthase kinase 3 beta) and *Nkg7* (Natural killer cell granule protein 7, a cytotoxic marker), and upregulation of *Myc, Jun* and *Il1b* (Interleukin 1 beta; Supplementary Table 2). Importantly, FOXO1, GSK3B and Myc are required in the maintenance of memory state of CD8^+^ T cells ^33, 34, 35^. Further analysis of DEGs revealed a significant enrichment of FOXO1 targets (n=35) ^36^ among 202 DEGs (Fig. 2j, *P value* = 1.2×10^6^), indicating a perturbation in FOXO1 signaling in CD8^+^ T cells from *Cxcr3^−/−^* mice. Taken together, our data suggested that metformin reprograms the CD8^+^ T cells phenotype and shapes the generation of the CD8^+^CXCR3^+^ T_M_ population. This generation of CD8^+^CXCR3^+^ T_M_ cells could be mediated by FOXO1, which can also control CD8^+^ T_M_ responses during infection ^37^.

### Metformin promotes mitochondrial spare respiratory capacity, mitochondrial health and survival of CD8^+^ T cells

Enrichment of CD8^+^ T_M_ cells by metformin suggested an effect dependent on regulation of cellular metabolism. Therefore, we measured the bioenergetic profiles of spleen CD8^+^ T cells from untreated and metformin-treated naïve WT mice. Oxygen consumption rate (OCR), an indicator of OXPHOS, was higher in metformin-treated CD8^+^ T cells compared to untreated CD8^+^ T cells (Fig. 3a). Importantly, metformin-treated CD8^+^ T cells demonstrated a larger mitochondrial SRC when compared to CD8^+^ T cells from untreated mice (Fig. 3a). This increase in SRC was found to be dependent on CXCR3 expression (Fig. 3b). The absence of metformin-mediated increased OXPHOS in CD8^+^ T cells from *Cxcr3^−/−^* mice could be due to the inherent reduced expression of *Foxo1* in these cells (Fig. 2i). Of note, FOXO1 regulates oxidative stress responses, proliferation, cell survival and apoptosis, and connects metabolic activity with immune response ^38, 39, 40^.

**Figure 3.**
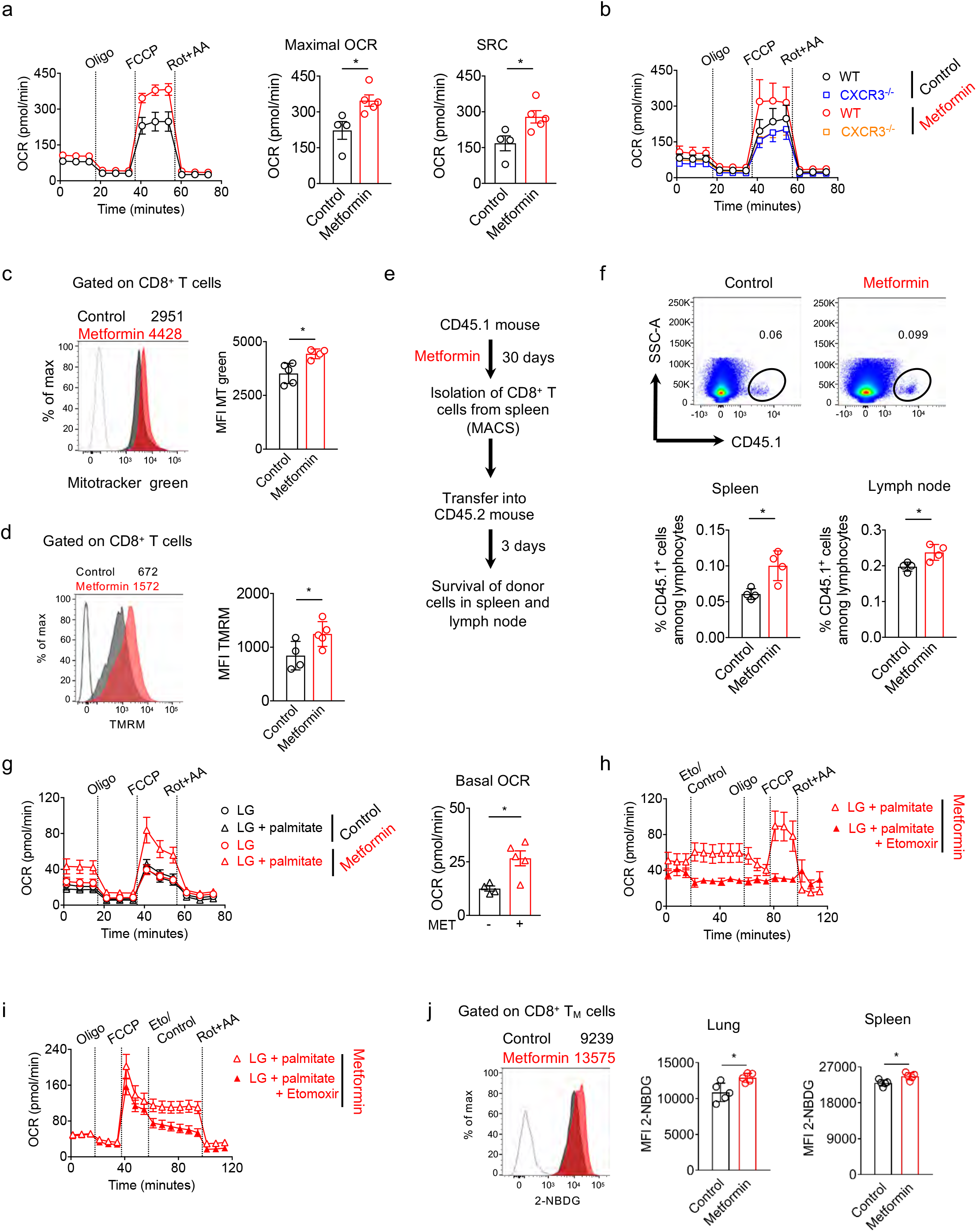
Metformin promotes CD8^+^ T cell spare respiratory capacity, mitochondrial mass, survival and FAO. **a** Oxygen consumption rate (OCR) and spare respiratory capacity (SRC) in spleen CD8^+^ T cells from control and metformin-treated WT mice in response to different drugs. **b** OCR in spleen CD8^+^ T cells from control and metformin-treated *CXCR3^−/−^* mice. **c** Mitotracker green staining of CD8^+^ T cells from control and metformin-treated WT mice. MFI - Median fluorescence intensity. **d** TMRM staining of CD8^+^ T cells from control and metformin-treated WT mice. **e** Experimental schematic of adoptive transfer of CD8^+^ T cells into congenic WT mice. **f** Frequency of CD45.1^+^ cells in the spleen and lymph node of recipient mice. **g** OCR in spleen CD8^+^ T cells from control and metformin-treated WT mice in response to different drugs under low glucose (LG) condition in the presence and absence of palmitate. **h, i** OCR in spleen CD8^+^ T cells from metformin-treated WT mice in response to different drugs under LG condition in the presence of palmitate. **j** Glucose uptake (2-NBDG) in CD8^+^ T_M_ cells, analysed by flow cytometry. Data shown as mean ± SD of 4 - 10 mice per group. \**p* ≤ 0.05, Mann-Whitney *U* test.

Enhanced SRC in cells might be explained by increased cellular mitochondrial mass ^20^. CD8^+^ T cells from metformin-treated WT mice showed increased Mitotracker Green median fluorescence intensity (MFI) compared to CD8^+^ T cells from untreated mice, indicating increased mitochondrial mass (Fig. 3c and Supplementary Fig. 5a). Metformin-educated CD8^+^ T cells from WT mice also showed a trend of increased mRNA expression for *Sell* (*CD62L*), *Cpt1a* and *Tfam* (mitochondrial transcription factor A) genes (Supplementary Fig. 5b). Additionally, metformin-educated CD8^+^ T cells showed increased accumulation of tetramethylrhodamine, methyl ester (TMRM), indicating healthy and functional mitochondria (Fig. 3d). In our experiments, we did not find evidence of an increased extracellular acidification rate (ECAR; indicator of glycolysis) in metformin-educated CD8^+^ T cells, when this was evaluated using glycolytic-rate assay (data not shown).

To further assess the impact of metformin on the CD8^+^ T_M_ compartment, we characterized the survival capacity of CD8^+^ T cells *in vivo* by adoptively transferring isolated metformin-treated or untreated cells into congenic mice (Fig. 3e). After 3 days, we found significantly more CD45.1^+^ donor metformin-treated CD8^+^ T cells in spleen and lymph node of the recipient mice (Fig. 3f). Together, our data suggested that metformin enhanced mitochondrial SRC and health of CD8^+^ T cells and promoted their survival, which are characteristic features of T_M_ cells.

### Metformin-educated CD8^+^ T cells use glucose for mitochondrial FAO and OXPHOS

Since metformin promoted the enrichment of the CD8^+^ T_M_ cells (Fig. 2) with increased SRC (Fig. 3a), we reasoned that metformin-educated CD8^+^ T cells might utilize extracellular glucose to fuel mitochondrial FAO and OXPHOS, a property of IL-15-induced CD8^+^ T_M_ cells ^20^. We evaluated this by measuring OCR in CD8^+^ T cells cultured in low glucose (LG) medium. The increased OCR and SRC seen in CD8^+^ T cells from metformin-treated mice cultured in glucose-sufficient medium (25mM, Fig. 3a) was not observed under LG conditions (0.5mM, Fig. 3g). This suggested that glucose was driving FAO and OXPHOS in metformin-educated CD8^+^ T cells. Additionally, under LG conditions, supplementation with fatty acid (FA; palmitate) rescued the SRC of metformin-educated CD8^+^ T cells (Fig. 3g), indicating that increased SRC of metformin-educated CD8^+^ T cells in glucose-sufficient media (Fig. 3a) was derived from FAO.

Next, we used the CPT1a inhibitor etomoxir (Eto), which blocks mitochondrial FAO, to confirm that the enhanced SRC in CD8^+^ T cells from metformin-treated mice depended on FAO. Treatment with Eto, either before or after carbonylcyanide-p-trifluoromethoxyphenylhydrazone (FCCP) injection, impaired basal OCR and SRC, respectively, in metformin-educated CD8^+^ T cells (Fig. 3h, i). Since CPT1a-independent effects of Eto on mitochondrial respiration have been observed ^41^ and we used higher concentration of Eto (viz. 200μM), FAO independent effect of metformin on OXPHOS cannot be ruled out. To further support the hypothesis that metformin-educated CD8^+^ T_M_ cells used glucose for FAO and OXPHOS, we performed glucose uptake assay using the fluorescent glucose analogue 2-NBDG. Lung and spleen CD8^+^ T_M_ cells from metformin-treated mice acquired more 2-NBDG than CD8^+^ T_M_ cells from untreated mice (Fig. 3j). We did not find an increased uptake of 2-NBDG by CD8^+^ T_E_ cells from metformin-treated mice (Supplementary Fig. 5c), indicating minimal effect of metformin on the metabolic reprogramming of those cells. Together, our data suggested that glucose fuels mitochondrial FAO which might be responsible for the induction of OXPHOS in metformin-educated CD8^+^ T cells.

### Distinct gene expression signatures in metformin-educated CD8^+^ T_M_ cells

We next used RNA sequencing to characterize the effect of metformin on splenic CD8^+^ T_M_ cells (Fig. 4a). A total of 267 genes was found to be differentially expressed in metformin-educated CD8^+^ T_M_ cells (Fig. 4a and Supplementary Table 4). IPA based molecular function enrichment revealed an effect of metformin on the expression of 86 genes associated with cell survival (Fig. 4b and Supplementary Table 5). Among these 86 genes we found an upregulation of *Cdk1* (cyklin dependent kinase 1) and *Agap2* (Arf-GAP with GTPase, ANK repeat and PH domain-containing protein 2) genes, consistent with a role of metformin in modulating CD8^+^ T_M_ cell fate (Fig. 4c). Additionally, metformin reduced the expression of *Mtor* (mammalian target of rapamycin), *Immt* (inner membrane mitochondrial protein, mitofilin), and *Casp1* (caspase-1) genes (Fig. 4c). Metformin is known to inhibit mTOR ^42^, while IMMT deficiency has been associated with perturbed mitochondrial function ^43^. Since caspase-1 is required for pyroptosis, a form of programmed cell death ^44^, we reasoned that the enhanced survival of metformin-educated CD8^+^ T cells (Fig. 3f) might be explained by the reduced caspase-1 activity. As predicted, metformin-treated *Mtb*-infected mice had reduced numbers of CD8^+^caspase-1^+^ cells in the spleen compared to the untreated *Mtb*-infected mice (Fig. 4d).

**Figure 4.**
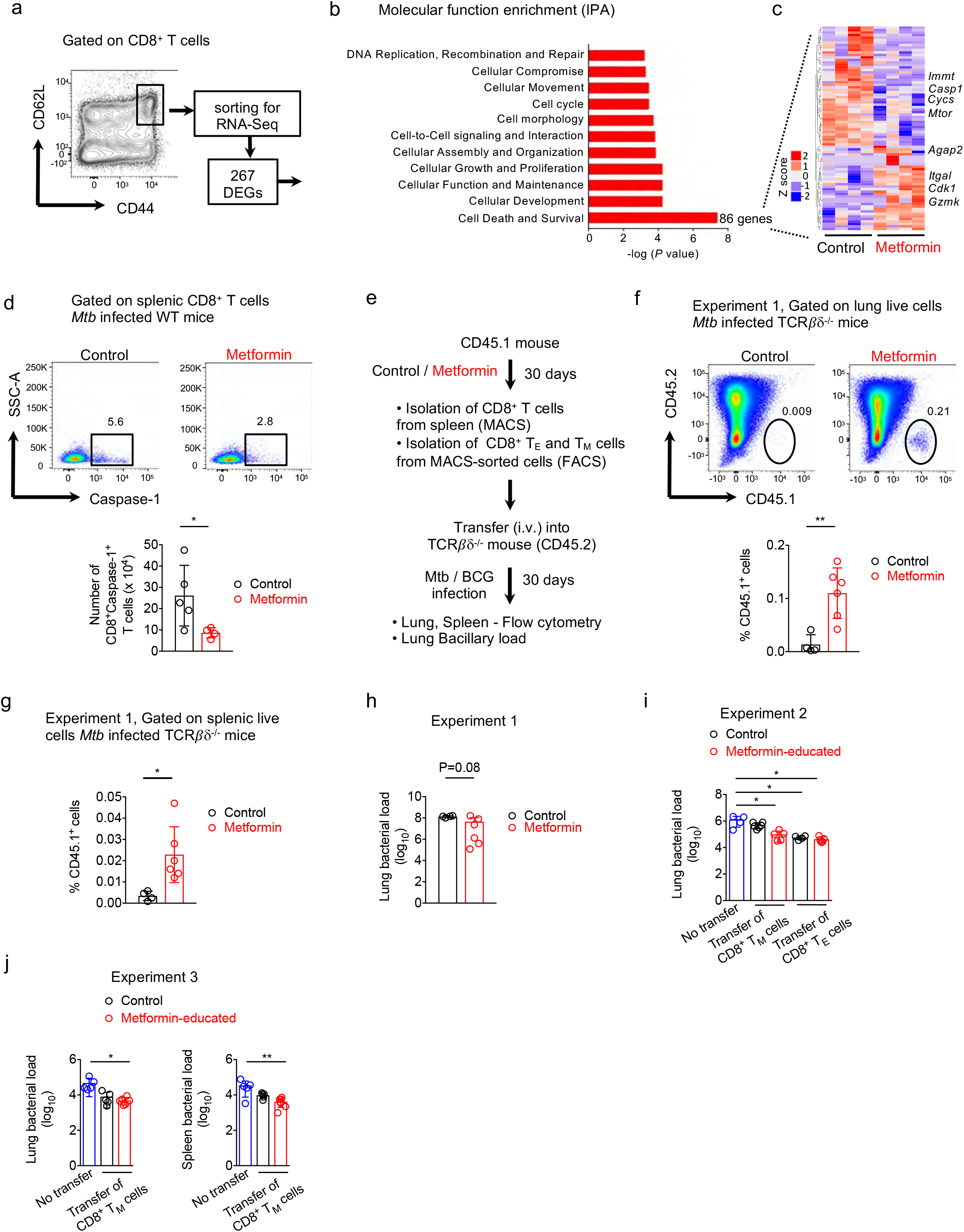
Metformin reprograms gene expression of CD8^+^ TM cells. **a** Schematic depicting sorting of CD8^+^ T_M_ cells for RNA sequencing and identification of DEGs in metformin-educated CD8^+^ T_M_ cells. **b** Molecular function enrichment via Ingenuity pathway analysis (IPA) of DEGs in (A). **c** Heat map displaying the expression level of 86 genes related to cell death and survival. **d** Flow cytometric analysis of caspase-1 in splenic CD8^+^ T cells from *Mtb*-infected mice treated or not with metformin. **e** Experimental set-up for the transfer of CD8^+^ T_M_ or T_E_ cells into *TCRβδ^−/−^* mice followed by *Mtb* or BCG infection. Three experiments were performed. **f, g** Flow cytometric analysis of the frequency of CD45.1^+^ lung (F) and splenic (G) cells in recipient mice. **h** Bacterial load in the lung of *Mtb*-infected *TCRβδ^−/−^* recipient mice. **i, j** Bacterial load in the lung of BCG-infected *TCRβδ^−/−^* recipient mice. Data of mean ± SD of 4 −5 mice per group, \**p* ≤ 0.05, \*\**p* ≤ 0.01, ANOVA with Tukey’s multiple comparison test. Data in d, f and g is mean ± SD of 4 −6 mice per group, \**p* ≤ 0.05, \*\**p* ≤ 0.01, Mann-Whitney *U* test.

To test whether metformin-educated CD8^+^ T_M_ cells survived better *in vivo* under infection settings, we adoptively transferred flow-sorted CD8^+^ T_M_ cells into congenic *TCRβδ^−/−^* mice that were subsequently infected with *Mtb* (Fig. 4e; total 3 experiments). We found significantly more metformin-educated donor cells in the lungs (Fig. 4f) and spleens (Fig. 4g) of the recipient *TCRβδ^−/−^* mice. This paralleled with a trend for lower bacterial load in the lungs of *TCRβδ^−/−^* mice that received metformin-educated CD8^+^ T_M_ cells compared to those receiving control CD8^+^ T_M_ cells (1.4 log10 reduction), although this did not reach statistical significance (Fig. 4h). In other experiments, *TCRβδ^−/−^* mice receiving metformin-educated CD8^+^ T_M_ cells had significantly lower lung and spleen bacterial load compared with the *TCRβδ^−/−^* mice that did not receive any T cells (0.9-1.3 log10 reduction; Fig. 4i, j). In this experiment, CD8^+^ T_E_ cells from control and metformin-treated mice equally restricted lung bacterial loads in the *TCRβδ^−/−^* recipient mice to a comparable level (Fig. 4i). Collectively, our data indicated that metformin reprogramed gene expression of CD8^+^ T_M_ cells, increasing their survival and anti-mycobacterial properties.

### Metformin enhances BCG vaccine immunogenicity and protective efficacy

The generation of T_M_ cells has important implications for vaccine efficacy. Since metformin-educated CD8^+^ T_M_ cells display distinct phenotypes and function associated with host protection, we evaluated the effect of metformin on the response to vaccination with Bacillus Calmette–Guérin (BCG) (Supplementary Fig. 6a). In BCG-vaccinated mice, metformin treatment was associated with an increased frequency of PPD (purified protein derivative of *Mtb*)-specific IFNγ-secreting splenic CD8^+^ T cells (Fig. 5a) but not CD4^+^ T cells (Supplementary Fig. 6b), compared with untreated BCG-vaccinated animals. Similar trend was observed in PPD-specific CD8^+^ T_M_ and T_E_ cells from metformin-treated BCG-vaccinated mice (Fig. 5a). We also observed an increased frequency of IFNγ^+^ total CD8^+^ cells (unstimulated) from metformin-treated BCG-vaccinated mice (Fig. 5a). CD8^+^ T cells from metformin-treated BCG-vaccinated mice also demonstrated an increased mitochondrial mass (Fig. 5b) and health (Fig. 5c) similar to that seen in unvaccinated animals (Fig. 3b, c). We next performed RNA sequencing on flow-sorted CD8^+^ T_M_ cells from metformin-treated and untreated BCG-vaccinated mice. Comparative analysis revealed 607 DEGs that were enriched for genes associated with OXPHOS (Fig. 5d and Supplementary Table 6, 7). A heat map of OXPHOS genes showed an increased expression of electron transport chain (ETC) genes in CD8^+^ T_M_ cells from metformin-treated BCG-vaccinated mice (Supplementary Fig. 6c).

**Figure 5.**
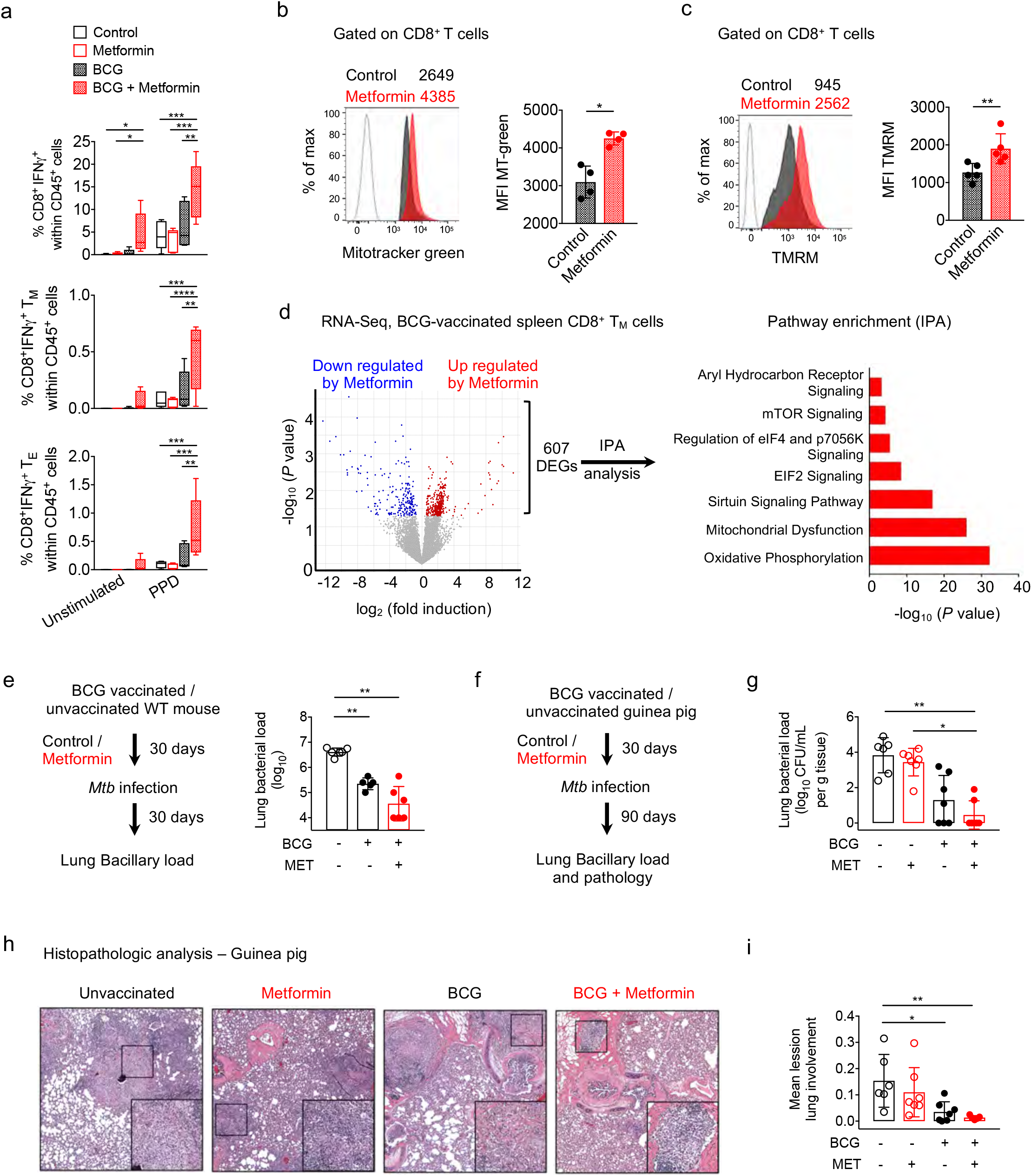
Metformin treatment promotes BCG immunogenicity and vaccine efficacy in mice and guinea pigs. **a** PPD specific response of spleen cells, analysed by flow cytometry. Data on IFNg producing CD8^+^ T, T_M_ and T_E_ cells from one out of two independent experiments is shown. Mean ± SD (n = 5). \**p* ≤ 0.05; \*\**p* ≤ 0.01; \*\*\**p* ≤ 0.001, ANOVA. **b, c** Mitotracker green (B) TMRM (C) staining of splenic CD8^+^ T cells from control and metformin-treated BCG-vaccinated WT mice. **d** Changes in gene expression in CD8^+^ T_M_ cells from BCG-vaccinated mice treated or not with metformin. IPA analysis of 607 DEGs. N=5 mice/group. **e** Schematic of *Mtb* infection of metformin-treated BCG-vaccinated mice. Lung bacillary load at 30 days post-infection (p.i) is shown. **f** Schematic of *Mtb* infection of metformin-treated BCG-vaccinated guinea pigs. **g** Lung bacillary load in BCG-vaccinated *Mtb*-infected guinea pigs treated or not with metformin. **h** Light micrographs of hematoxylin and eosin (H&E) staining of lung sections from animals in g. Magnification for main image 40x and insert 100x. **i** Morphometric analysis of lung sections shown in h indicating the mean lesion lung involvement from each guinea pig. Data in b and c, is representative from two experiments. Data is indicated as means ± SD of 4 - 8 animals/ group, \**p* ≤ 0.05, \*\**p* ≤ 0.01, Mann-Whitney *U* test.

We next examined whether metformin treatment would enhance BCG vaccine-mediated *Mtb* control in mice. A trend for reduced *Mtb* bacillary load in the lungs of metformin-treated vaccinated mice compared to untreated vaccinated mice was observed (Fig. 5e). BCG-vaccinated metformin-treated mice showed an increased frequency memory CD8^+^ T cells, and decreased effector CD8^+^ T cells upon *Mtb* infection compared to the BCG-vaccinated mice alone (Supplementary Fig. 6d, upper panel). When stimulated with class I-restricted peptide (TB10.4) of the *Mtb EsxH* gene, memory CD8^+^ T cells from BCG-vaccinated metformin-treated mice demonstrated increased TNFa production compared to the BCG-vaccinated mice alone (Supplementary Fig. 6d, lower panel). We next used the guinea pig TB model to further evaluate metformin-mediated enhancement of BCG efficacy (Fig. 5f) ^45^. Lung *Mtb* bacillary load in metformin-treated BCG-vaccinated guinea pigs was significantly lower compared to unvaccinated animals or animals which received only metformin, reduction of 3.4 log10 and 2.9 log10 respectively (Fig. 5g). In this experiment, bacillary load in untreated vaccinated animals was not significantly different from unvaccinated animals (reduction of 0.4 log10, Fig. 5g). Histopathological evaluation of the infected lungs showed reduced infiltrating leukocytes and lesions in BCG-vaccinated guinea pigs compared with unvaccinated animals (4.3 fold reduction in lung lesion burden, Fig. 5h). Importantly, lungs of metformin-treated BCG-vaccinated guinea pigs showed 11.2 and 2.6 fold reduced lesion burden compared with the control and untreated BCG-vaccinated guinea pigs, respectively (Fig. 5h, i). Taken together, our data suggested that metformin enhances the BCG vaccine response and efficacy against *Mtb*.

### Metformin expands CD8^+^CXCR3^+^ T_M_ cells in humans

To investigate the effect of metformin in humans, we performed CyTOF on peripheral blood mononuclear cells (PBMCs) from healthy human participants who took a standard dose of metformin for five days (Fig. 6a, See methods) ^46^. The PBMCs were stained with a panel of 38 markers (Supplementary Table 8). Analysis of CD8^+^ T cells (CD45^+^CD14^−^CD19^−^LA-DR+CD3+CD4^−^γδTCR^−^TCRVα7.2^−^CD161^−^CD56’CD8^+^) using t-SNE (Supplementary Fig. 7a) identified T_N_ cells (CCR7^+^CD45RO^−^CD45RA^+^), T_E_ cells (CCR7^−^CD45RO^+^CD45RA^+^) and T_M_ cells (CCR7^+^CD45RO^+^CD45RA^−^) (Fig. 6b, c; Supplementary Fig. 7a) ^47^. Two distinct T_M_ and T_E_ subgroups were observed (Fig. 6b). Interestingly, the T_M2_ subset was positive for CD127 and CXCR3 (Fig. 6d), similar to what we observed in mouse CD8^+^ T_M_ cells (Fig. 2c). This CD8^+^ T_M2_ (CD8^+^CD127^+^CXCR3^+^) population expanded upon metformin intake in healthy subjects (Fig. 6e), and also showed increased expression of anti-apoptotic BCL2 (Fig. 6f), which was previously shown to correlate with CD8^+^ T cell survival ^48^. We next performed flow cytometric analysis on the PBMCs from T2D patients who were being treated with either metformin or with other antidiabetic drugs as monotherapy (Supplementary Fig. 7b and Supplementary Table 9). These two groups had similar fasting plasma glucose levels (Supplementary Table 9). Metformin-receiving T2D patients showed an increased frequency of CD8^+^CXCR3^+^ T_M_ cells compared to T2D patients treated with other anti-diabetic drugs (Fig. 6g). Overall, our data indicated that metformin induced the expansion of memory-like CD8^+^CXCR3^+^ T cells both in mice and humans.

**Figure 6.**
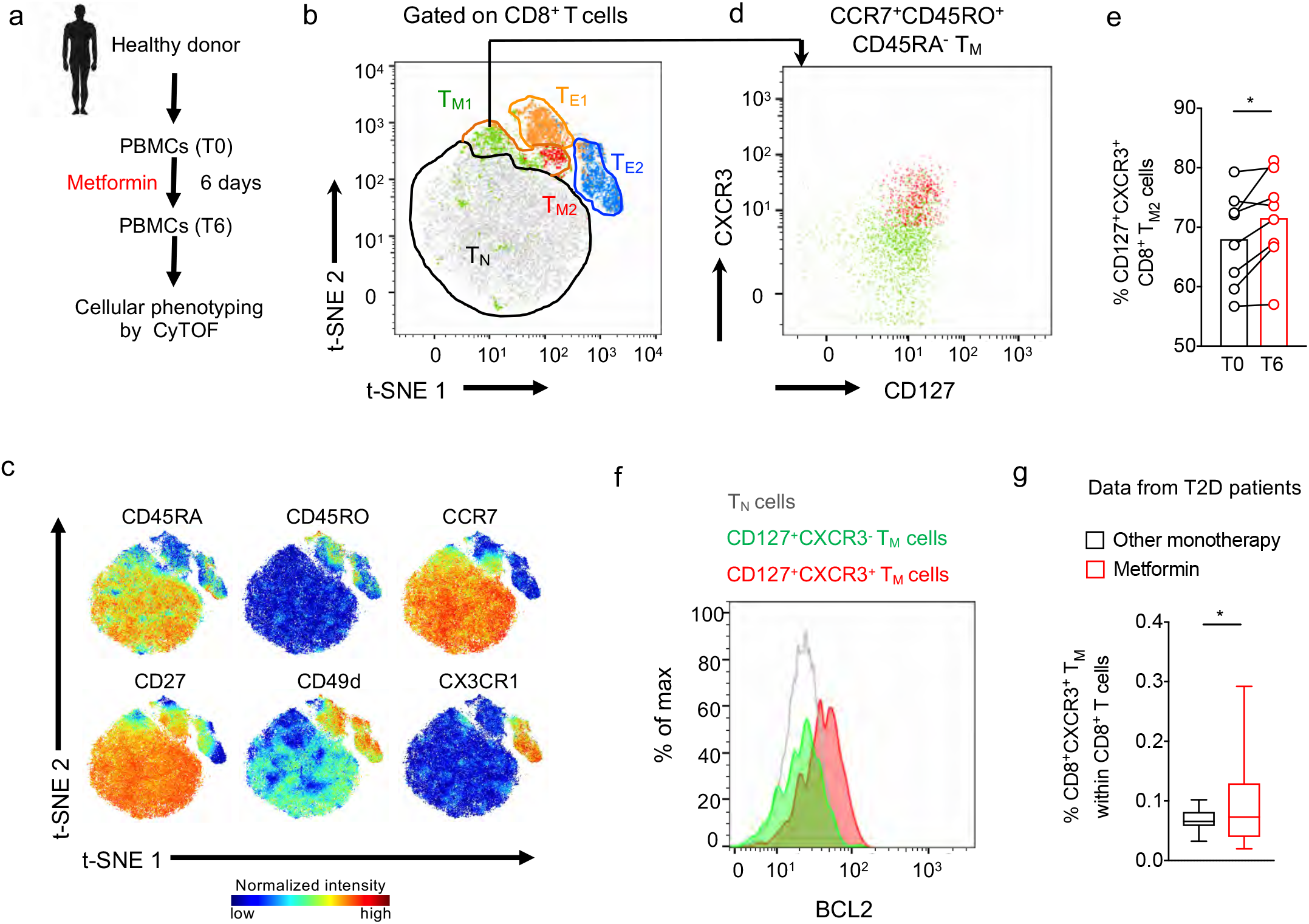
Metformin treatment expands CD8^+^CXCR3^+^ TM cells in healthy and diabetic individuals. **a** Strategy of analysing PBMCs from metformin treated healthy individuals. **b** t-SNE map of human peripheral CD8^+^ T cells with manually cluster gating for T_E_, T_M_ and T_N_ cells. **c** Normalized expression intensities of indicated markers were calculated and overlaid on the t-SNE plot, characterising all cluster of human CD8^+^ T cells. **d** Manual gating identified differential expression of CD127 and CXCR3, distinguishing two CD8^+^ T_M_ subpopulations, T_M1_ and T_M2_. **e** Frequency of CD8^+^CD127^+^CXCR3^+^ T_M_ cells before and after metformin treatment. N=8, \**p* ≤ 0.05, Paired Wilcoxon test. **f** Overlay plot showing expression of BCL-2 in CD8^+^CD127^+^CXCR3^−^ and CD8^+^CD127^+^CXCR3^+^ T_M_ cells. BCL2 expression in CD8^+^ T_N_ cells is also indicated. Data shown from one representative donor. **g** Flow cytometric analysis of frequency of peripheral CD8^+^CXCR3^+^ T_M_ cells in DM patients receiving either metformin or other diabetic monotherapy, mean ± SD of 18 - 42 patients/group, \**p* ≤ 0.05, Mann-Whitney *U* test.

### Effect of metformin on Mtb clearance and reactivation

CD8^+^ T cells play an important role in long-term protective immune responses against *Mtb* infection and reactivation of latent TB ^10, 49^. To assess the persistence of metformin effects, we first measured the bioenergetic profiles of splenic CD8^+^ T cells from naïve mice that received metformin for 6 months. Similar to our previous finding (Fig. 3a), OCR was higher in CD8^+^ T cells from mice chronically treated with metformin compared to those from untreated mice (Supplementary Fig. 8a). This was associated with an increased number of splenic CD8^+^ T_M_ cells in metformin treated mice (Supplementary Fig. 8b). We next used a modified version of the TB Cornell model ^50^ to evaluate the long-term effects of metformin on *Mtb* clearance and reactivation. In this model, *Mtb*-infected mice were treated with pyrazinamide (PZA) and isoniazid (INH) ± metformin for 70 days starting on day 40 p.i., followed by a rest period of 40 days, and then challenge with dexamethasone to reactivate any persisting bacilli (Supplementary Fig. 8c). In the animals which received metformin along with PZA and INH bacilli was recovered from the lungs of 40-50% mice sacrificed 40 days after the drugs were stopped (day 150 p.i, Table 1 and Supplementary Fig. 8c). However for mice which received only PZA and INH, bacilli were recovered from 75-80% of animals at day 150 p.i. After 15 days of dexamethasone treatment bacilli were recovered from the lungs of 20-50% of mice treated with metformin+PZA+INH compared to 75-80% mice treated with PZA+INH (Table 1). Taken together these data suggest that metformin enhances the sterilizing activity of suboptimal antimicrobial treatment for *Mtb* infection by boosting intrinsic host immune function.

**Table 1.** Efficacy of metformin against *Mtb* in modified Cornell-mice Model*.

| **EXP No.** | **Day 150** |  | **Day 167** |  |
| --- | --- | --- | --- | --- |
|  | **PZA+INH‡** | **MET+PZA+INH§** | **PZA+INH** | **MET+PZA+INH** |
| Expt 1 | 1/5 (20%) | 3/5 (60%) | 1/5 (20%) | 4/5 (80%) |
| Expt 2 | 1/4 (25%) | 2/4 (50%) | 1/4 (25%) | 2/4 (50%) |

*Definition of abbreviations:* INH= isoniazid; PZA= Pyrazinamide; MET = Metformin.
\* Proportion of mice with no detectable CFUs in the lungs and spleen during PZA+INH therapy with and without MET in two different experiments.
‡*Mtb*-infected mice were administered INH (10 µg/mL) and PZA (15 mg/mL)
<sup>||</sup>Number of mouse with no detectable CFU out of total number of animals in that group. In bracket percentage information is provided.

## Discussion

The goals of HDT for TB are to accelerate lesion sterilization and mitigate immune pathology by educating host anti-*Mtb* responses ^2, 51^. Previously, we proposed metformin as an adjunctive TB treatment, reporting that it lowers risk for cavitary TB and mortality in adults with T2D ^3^. In the present study, we showed that metformin treatment (i) expands CD8^+^CXCR3^+^ T_M_ cells in mice and humans, (ii) reprograms CD8^+^ T cells metabolic and transcriptional circuits, (iii) enhances BCG vaccine protective efficacy, and (iv) improves sterilizing ability of antibiotics thus preventing *Mtb* reactivation.

Metformin-educated Ag-inexperienced CD8^+^CXCR3^+^ T_M_ cells share phenotypic and functional properties with Ag-specific CD8^+^ T_CM_, Ag-inexperienced T_VM_, innate-memory CD8^+^ T, and/or IL-15-derived CD8^+^ T_M_ cells ^52^, and have enhanced survival and anti-*Mtb* properties. Cytokine-activated CD8^+^ T_M_ cells are known to contribute to host protection against diverse pathogens ^17^. The differential transcriptomic and phenotypic properties of metformin-educated CD8^+^CXCR3^+^ T_M_ cells suggest that these cells may represent another type of “innate CD8^+^ T cell sensor” ^17^. The CXCR3/CXCL10 axis can shape the distribution of CD8^+^ T_M_ cells in the tissues of non-immunized mice ^53^ and CXCR3-expressing innate CD8^+^ T_M_ cells are known to express activation markers, respond to IL-2 and IL-15, and possess antibacterial effector function ^54, 55^. Additionally, CXCR3 is required for the migration and establishment of CD8^+^ T_CM_ cell population in the respiratory tract, where they protect against influenza A virus infection ^29^.

CD8^+^ T_M_ cells protect the host against infectious diseases by virtue of their recall ability to expand and execute effector functions ^10, 56^. These responses are promoted by AMP-activated protein kinase (AMPK) ^57^, a crucial energy sensor and an indirect target of metformin. AMPK signaling promotes OXPHOS during CD8^+^ T_M_ generation ^58^ and maintains mitochondrial health in T_M_ cells ^59^. This indicates the importance of AMPK and OXPHOS in rewiring of metformin-educated Ag-inexperienced CD8^+^ T_M_ cells in our experiments. We demonstrated that CXCR3 signaling is required for metformin-mediated induction of OXPHOS in CD8^+^ T cells, suggesting a possible CXCR3- and AMPK-dependent effect(s). How CXCR3 regulates metformin-mediated induction of OXPHOS, mitochondrial health and expansion of CD8 memory remains to be determined. Interestingly, mice devoid of AMPK in CD4^+^ T cells shows deficiency in the CXCR3 expression ^60^. Our transcriptomic analysis of CD8^+^ T cells from *Cxcr3*^−/−^ mice suggested dependency of metformin-mediated reprograming of mitochondrial function on Foxo1 network (Fig. 2i. j), which is an important metabolic check point ^40^. Of note, TB immunopathology could be modulated by targeting T cells mitochondrial metabolism ^61^.

The recall ability of T_M_ cells is also the basis for vaccinations against numerous diseases including TB. The current TB vaccine, BCG, is unable to protect against pulmonary TB in adults, hence there is an urgent need to improve BCG or to develop new TB vaccine. Here we showed that metformin-treatment resulted in an increased (i) elicitation of Ag-specific CD8^+^ T_CM_ in the tissues of BCG-vaccinated mice, (ii) TNFa secreting Ag-specific CD8^+^ T_CM_ cells in the tissues of BCG-vaccinated *Mtb*-challenged mice, and (iii) protection of BCG-immunized mice and guinea pig against *Mtb* challenge. Importantly, expression of CXCR3 by TB vaccine-specific CD8^+^ T cells is needed for their homing to the lung mucosa ^62^. This raises the possibility of the presence of metformin-educated CD8^+^ T cells in different anatomical location within lung parenchyma, which is responsible for enhanced protective immunity. We did not see enhanced immunogenicity among Ag-specific CD4^+^ T_CM_ cells from metformin-treated mice, which goes in line with no benefit of metformin-educated Ag-inexperienced CD4^+^ T cells on *Mtb* growth in adoptive transfer experiments. Furthermore, these Ag-inexperienced CD4^+^ T cells also did not show increased

OXPHOS (data not shown). The differential effect of metformin on the metabolism of CD4^+^ and CD8^+^ cells in naïve or vaccinated host is not clear at present. However, since metformin normalizes CD4^+^ T cell mitochondrial function to alleviate lupus in B6.Sle1.Sle2.Sle3 mice ^63^ and aging-associated inflammation ^64^ in humans, an effect of metformin on CD4^+^ T cell metabolism in *Mtb*-infected hosts cannot be excluded. Nevertheless, our results support the concept of enhancing the efficacy of BCG vaccine by small molecule drugs ^65^.

In conclusion the data presented here show that metformin shifts CD8^+^ T cell metabolism, leading to the expansion of a memory-like CD8^+^CXCR3^+^ phenotype with functional properties that contribute to host control of *Mtb* infection and disease. Our findings are in line with evidence that metabolic reprograming by metformin reverses CD8^+^ T cell dysfunction during chronic *Mtb* infection ^66^. Our current and previously reported studies show that metformin ameliorates TB immune pathology, while others reported that it reverses established pulmonary fibrosis in the bleomycin model. That evidence, together with our demonstration that metformin enhances BCG vaccine immunogenicity and protective efficacy adds to the basis in evidence supporting metformin as a candidate for TB-HDT and as a TB vaccine adjunct.

## Methods

### Healthy human samples

Blood PBMCs from 8 male healthy non-obese volunteers without medication, kidney function loss or metabolic disorders, who received metformin for 5 days in increasing doses starting at 500 mg once a day and ending with 1000 mg twice a day were utilized ^46^. Blood was drawn at baseline time point before metformin intake (T0) and immediately after metformin intake on day 6 (T6).

### Diabetic human patient cohort

T2D patients included in the study are part of the Singapore Longitudinal Aging study 2 (SLAS-2), which is a population-based cohort intended to study the biology of aging among Singaporean elderly individuals above the age of 55 years old ^67^. PBMCs from the blood samples collected from overnight fasting individuals were washed with PBS and cryopreserved in liquid nitrogen. PBMCs were thawed and used for flow cytometric staining. Fasting blood glucose, blood count, hematocrit level, albumin, creatinine, estimated glomerular filtration rate was performed on the blood samples.

### Animals

C57BL/6J, CD45.1, *Cxcr3^−/−^, TCRβδ^−/−^* and OT-I mice were obtained from SIgN mouse core. CD45.1 and CD45.2 mice for chimeric *in vivo* experiments were obtained from Jackson Labs (Bar Harbor, USA), and performed at UMMS. For all *in vivo* experiments female C57BL/6J mice aged between 8 and 12 weeks were used. Hartley strain guinea pigs (350 – 400 g) were obtained from Charles Rivers (Wilmington, MA), maintained on a normal chow diet, acclimated for 1 week then randomly assigned to 3 treatment groups.

### Metformin treatment for mice and guinea pigs

Metformin (1,1-Dimethylbiguanide hydrochloride, Sigma Aldrich) was provided to mice in drinking water (1.83 mg/ml which is equivalent to 270 mg/kg). Details of metformin treatment for different experimental models is provided in the figures. Guinea pigs were treated with 25 mg/kg of metformin suspended in carrier (Ora-Sweet, Paddock Laboratories, Minneapolis, MN), or carrier alone, and administered orally once daily, for 30 days, prior to *Mtb* infection. The metformin dose was determined in guinea pigs based on PK/PD analysis in which the peak plasma concentration reached 1.1 μg/mL (peak plasma concentration in human 1.1 μg/mL).

### BCG vaccination

Female mice aged 8 - 10 weeks were vaccinated subcutaneous with a single dose BCG Pasteur strain (1 × 10^6^ colony-forming units (CFU) in 100 μl PBS). In BCG vaccinated mice metformin treatment was initiated 1 day post-vaccination. Guinea pigs were vaccinated intra-dermally (i.d) with 1 × 10^5^ BCG Pasteur strain. In vaccinated guinea pigs metformin treatment was initiated 14 days post-vaccination. Sham vaccinated guinea pigs received an equal volume of saline i.d.

### *Mtb* infection

Female mice were infected via aerosol or intra tracheal (i.t) ^68^ route with *Mtb* H37Rv or *Mtb* Erdman to deliver 100 - 200 viable bacilli per lung. For BCG infection experiments, 10 × 10^6^ BCG bacilli were used. The implanted bacterial loads in the lungs of infected mice were quantified one day post-infection. Guinea pigs were infected with the *Mtb* H37Rv by aerosol to deliver 10 - 20 bacilli per animal and confirmed by culture.

### Modified Cornell model

Starting 40 days p.i *Mtb*-infected female mice were given isoniazid (10 μg/mL) and pyrazinamide (15 mg/mL) in drinking water ad libitum ^69^ for 70 days. This was followed by a 40 day rest for all treatments. Subsequently, at day 150 p.i mice were treated with 1 mg/mL of dexamethasone in drinking water for 15 days.

### Culturing of mycobacterial strains

*Mtb* H37Rv, Erdman and *M. bovis* BCG were grown in Middlebrook 7H9 broth (BBL Microbiology Systems) supplemented with 10 % ADS (Albumin, Dextrose, Saline, Sigma) enrichment medium with 140 mM NaCl, 0.5 % glycerol and 0.5 % Tween-80 at 37°C for 5 to 10 days to an OD_600_ of 0.4 - 0.5. Bacterial cultures were harvested, re-suspended in 7H9 broth with 20% glycerol, and aliquots were frozen at −80°C until later use. Before infection, cells were thawed, washed and sonicated. viable cell counts per mL in thawed aliquots were determined by plating serial dilutions on to Middlebrook 7H11 agar plates supplemented with 10% OADC (Oleic acid, Albumin, Dextrose, Catalase, BBL Microbiology Systems) and 0.5 % glycerol followed by incubation at 37°C.

### Enumeration of mycobacterial CFU in the tissues of infected mice and guinea pigs

Mycobacterial loads in lung and spleen of euthanized mice were evaluated at different time points post infection by homogenizing the organs in PBS or PBS with 0.05 % Tween-80 using MACS tissue dissociator (Miltenyi Biotec). Guinea pigs were euthanized with an intra-peritoneal overdose of pentobarbital (>120 mg/kg), and representative lung tissue samples were collected sterilely and homogenized in 9 ml of sterile saline. Tenfold serial dilutions of tissue homogenates in sterile PBS were plated in triplicates on Middlebrook 7H11 agar plates supplemented with 10 % OADC and 0.5 % glycerol, incubated for 3-6 weeks at 37°C and CFUs counted.

### Cell isolation from lungs and spleens

Lungs were aseptically isolated from mice and incubated at 37°C in 10% FBS RPMI media containing 1.4 mg/ml collagenase A (Roche) and 30 μg/ml DNase I for 60 min. The treated lung tissue and spleen was dissociated over the 70 μm cell strainer (Fisherbrand). Strainer was washed to collect single cell suspension. RBC were lysed with ACK lysing buffer (Lonza) for 5 min following by washing of cells with culture media. Cells were counted and adjusted to 5 × 10^6^ cells/ml.

### Magnetic sorting of murine T cells

Spleen cells were washed twice in fresh MACS buffer. CD4^+^ and CD8^+^ T cells were magnetically purified via negative selection using Dynabeads or MACS cell separation system (Supplementary Table 10) according to the manufacturer’s protocols, followed by purity check which was around 90-95%.

### FACS sorting of murine memory CD8^+^ T cells

MACS-sorted splenic CD8^+^ T cells were stained with live dead marker, blocked with Fc block and surface stained with anti-CD3-FITC, anti-CD4-Pacific blue, anti-CD8-APC-Cy7, anti-CD44-BV650 and anti-CD62L-PE-Cy7 (Supplementary Table 11). Stained cells were sorted on FACS Aria II, FACS Aria III and BD Influx (Becton Dickinson, San Jose, CA, USA), followed by purity check which was around 90-95%.

### Adoptive T cell transfers

Purified splenic CD4^+^ or CD8^+^ T cells from untreated or metformin-treated CD45.1 mice were transferred via tail vein injection into sub lethally irradiated (600 rads) CD45.2 WT mice (Fig. 1). MACS-sorted splenic CD8^+^ T cells or T_M_ or T_E_ from untreated or metformin-treated CD45.1 mice were transferred via intra orbital injection into CD45.2 WT or *Tcrβδ^−/−^* (Fig. 3d, e). The details of number of T cells transferred is in Supplementary Table 12.

### Metabolism assays

Real-time mitochondrial OCR and ECAR of splenic CD8^+^ T cells were measured in media (DMEM containing 25 mM glucose and 2 mM L-glutamine), in response to 1 μM oligomycin, 5 μM FCCP and 0.5 μM rotenone/antimycin A, using Seahorse XF Cell Mito Stress and Glyco Stress Test Kit and XFe96 Extracellular Flux Analyzer (Agilent Technologies, USA, Supplemental Table 8). FAO assay were carried out using Seahorse XF Palmitate-BSA FAO Substrate kit in media (KHB containing 2.5 mM glucose and 0.5 mM Carnitine) in response to 1 μM oligomycin, 1 μM FCCP and 0.5 μM rotenone/antimycin A. For FAO assay, cells were prior starved in Seahorse DMEM medium (DMEM containing 0.5 mM glucose, 1 mM L-glutamine and 0.5 mM Carnitine) for 2 hours. Also for FAO assay, BSA or Palmitate:BSA was added into the 96-well plate and 200 μM etomoxir were used additionally as drug injection. SRC was determined as the absolute increase in OCR after FCCP injection compared with basal OCR. Data was analysed with Wave software, version 2.4 (Agilent Technologies).

### Flow cytometry

Single cell suspensions of spleens, lungs and lymph nodes (from mice) were prepared at the indicated time-points. Live dead staining was performed for 15 minutes at room temperature or for 20 minutes at 37°C. Non-specific antibody binding was blocked by adding anti-CD16/32 (Fc-block) (BD Biosciences). Following this surface markers were stained for 20-25 minutes at 4°C with the following anti-mouse antibodies: anti-CD45-Pacific blue/BV605/ FITC, CD45.1-BV711, CD45.2-APC, anti-CD3-FITC/ Pacific blue/ BUV395/ PerCP-Cy5.5, anti-CD4-Pacific blue/ Alexa Fluor 700/PerCP-Cy5.5/PE, anti-CD8-APC-Cy7/ FITC, anti-CD44-BV650/ PE-Cy7/ PE-Cy5, anti-CD62L-PE-Cy7/ PE-CF594 and anti-CXCR3-BV605 (Supplementary Table 11). In some experiments, mitochondrial staining was performed with 200 nM Mitotracker green and/or 20 μM Tetramethylrhodamine, methyl ester (TMRM) in RPMI media without FCS for 30 minutes at 37°C prior to staining of surface markers.

For intracellular cytokine staining, spleen cells were *ex vivo* stimulated for 20 h with 40 μg/ml PPD (purified protein derivative, Statens Serum Institute, Denmark) in the presence of Golgi Stop™ (BD Biosciences) and Monensin (1000x, BioLegend) for the last 4 hours, at 37°C and 5 % CO2. In some experiment’s spleen cells were *ex vivo* stimulated with IL-2 and TB10.4 peptides for 5 hours at 37°C. Brefeldin A was added to the stimulation for the last 4 hours. Live dead staining and blocking was performed. Cells were then stained with anti-CD45-BV605, anti-CD3-Pacific blue/ PerCP-Cy5.5, anti-CD4-PE/ Alexa Fluor 700, anti-CD8a-APC-Cy7/FITC, anti-CD44-PE-Cy7/ PE-Cy5 and anti-CD62L-AlexaFluor 700/ PE-Cy7 antibodies for 25 min at 4°C (table S9). After washing, cells were fixed and permeabilized for 20 minutes with Cytofix/ Cytoperm™ (BD Biosciences). Permeabilized cells were washed with Perm/Wash buffer™ (BD Biosciences) and stained with anti-IFN-γ-APC/ BV 421, TNF-a-APC or Perforin-PE antibody for 25 minutes at 4°C. Subsequently, cells were fixed in 2 % paraformaldehyde before acquisition.

Human PBMCs were stained with live dead marker and following anti-human antibodies: anti-CD34-PE, anti-CD4-BV750, anti-CD8-BV605, anti-CD14-BV605, anti-TCRγ/δ-APCR700, anti-CD27-BV510, anti-CD45-BUV805, anti-CxCR3-BV650, anti-TCRVδ2-BV711, anti-CD3-BV570, anti-Vα7.2-APC-Cy7, anti-CD45RO-PerCP/Cy5.5, and anti-CD161-BV785 (Supplementary Table 13). Stained cells were acquired using a LSRII cytometer with 5 lasers (BD, San Jose, CA, USA) or Fortessa cytometer with 4 lasers (Becton Dickinson, San Jose, CA, USA) or Symphony cytometer (BD Biosciences, California, USA). Flow data was analysed with FlowJo software (Tree Star, Inc., USA).

### Assessment of caspase-1 activity

The activity of caspase-1 was determined using fluorescent inhibitor of caspase-1 (FAM-Flica Caspase-1 Assay kit, ImmunoChemistry Technologies). Staining procedure was performed according to manufactures protocol. Briefly, after live/dead and surface antibody staining of lung and spleen single cells, FLICA reagent FAM-YVAD-FMK was provided to single cells and binding to activated caspase-1 was allowed for 1 hour at 37°C in the dark. Stained cells were washed with apoptosis wash buffer, followed by fixation. The green fluorescent was detected in FITC channel and samples were acquired using LSRII cytometer.

### Glucose uptake (2-NBDG) assay

2-(N-(7-nitrobenz-2-oxa-1,3-diazol-4-yl)amino)-2-deoxyglucose (2-NBDG) uptake was performed on glucose-deprived cultures of spleen and lung single cells. Cells were incubated in media containing 200 μM 2-NBDG for 20 minutes at 37°C. Followed this live dead and surface marker staining was performed. Fluorescent of 2-NBDG was detected in FITC channel using Fortessa cytometer.

### CyTOF staining and data analysis

Spleen and lung cells from mice, and human PBMCs were first stained with 200 μM cisplatin (Sigma) for the discrimination of dead cells. Upon this, cells were washed and stained with a primary antibodies cocktail for 30 minutes at 4°C (Supplementary Table 1 (for human antibodies) and Supplementary Table 8 (for mouse antibodies)). After washing, cells were stained with the heavy metal-labelled antibodies cocktail for 30 minutes at 4°C. Cells were washed, fixed and permeabilised using fixation FoxP3 buffer (eBioscience) for 45 minutes at 4°C. Upon this cells were stained with intranuclear antibodies cocktail for 30 minutes at 4°C. Labelled cells were washed twice with PBS and fixed in 2 or 4 % PFA overnight. Subsequently, cells were washed and stained for barcodes (30 minutes at 4°C) and DNA (10 minutes at 4°C). Samples were acquired on the Helios CyTOF (Fluidigm), and analysed as mentioned before ^24, 25, 70^. For details see supplemental methods.

### *Ex vivo* culture of primary T cells

OT-1 splenocytes were activated with 20 μg/ml OVA protein (Invivogen) and 100 U/ml hIL-2 (Miltenyi Biotec) and cultured in IMDM media (supplemented with 10 % FBS, 2 mM L-glutamine, 1 mM Sodium pyruvate, 0.1 mM NEAA, 1 % Kanamycin) for 3 days. After washing, cells were subsequently cultured with 0.4, 2, 10 ng/ml of hIL-15 (R&D) and/or 2 mM MET for 4 days. IL-15-derived antigen-specific CD8^+^ T cells were analysed for memory markers by flow cytometry.

### Real-time quantitative RT-PCR

Cells or tissues were lysed in TRIzol Reagent (Invitrogen), and total RNA was extracted using the RNeasy Mini Kit (Qiagen). RNA was reverse-transcribed using the iScript cDNA synthesis Kit (Bio-Rad). Complementary DNA (cDNA) was used for real-time quantitative PCR using 2x iQ SYBR Green supermix and C1000 Thermal Cycler (Bio-Rad) using specific primers (see supplemental methods). The relative mRNA expression level of each gene was normalized to *Gapdh* housekeeping gene expression in the untreated condition, and fold induction was calculated by the ΔΔCT method relative to those in untreated cells per tissue.

### RNA sequencing and analysis

Total RNA was extracted following the double extraction protocol: RNA isolation by TRIzol followed by a Qiagen RNeasy Micro clean-up procedure (Qiagen, Hilden, Germany). The extracted RNA samples were inspected on Agilent Bioanalyzer (Agilent Technologies, Santa Clara, CA) for quality assessment. All RNA samples were analysed on Perkin Elmer Labchip GX system (Perkin Elmer, Waltham, MA, USA) for quality assessment with a median RNA Quality Score of 9. cDNA libraries were prepared using 500 pg of total RNA and 1ul of a 1:200,000 dilution of ERCC RNA Spike in Controls (Ambion, Thermo Fisher Scientific, Waltham, MA, USA) using SMARTSeq v2 protocol ^71^ except for the following modifications: 1. Use of 20 μM TSO; 2. Use of 250 pg of cDNA with 1/5 reaction of Illumina Nextera XT kit (Illumina, San Diego, CA, USA). The length distribution of the cDNA libraries was monitored using DNA High Sensitivity Reagent Kit on the Perkin Elmer Labchip GX system (PerkinElmer, Waltham, MA, USA). The libraries were qPCR quantified (Kapa Biosystems, Wilmington, MA) to ascertain the loading concentration. RNA Seq libraries of FACS sorted CD8 T_M_ cells subjected to an indexed PE sequencing run of 2 × 51 cycles on an Illumina HiSeq 2000. FASTQ files were checked for quality control issues using fastQC and none were found. The reads in the FASTQ files were mapped onto the mouse genome build mm10 using STAR and the gene counts counted using featureCounts (part of the Subread package). Differentially expressed genes (DEGs) were detected using DESeq2 running in R version 3.3.3. DEGs having a p value less than 0.05 were considered to be statistically significant. DEGs were used in Ingenuity Pathway Analysis (IPA) for a core analysis. Heat maps were generated using the package ComplexHeatmap in R version 3.3.3.

### Histology and morphometry of guinea pigs lung samples

Left caudal lung lobe of guinea pigs was collected for histopathology and was fixed in 4 % paraformaldehyde for >48 hours, processed and embedded in paraffin, sectioned and stained with hematoxylin and eosin (H&E). The *Mtb* lesion burden was determined morphometrically using the area fraction fractionator method with Stereo investigator software (MBF Biosciences, Williston, VT). The data was expressed as the percent of lung area affected by characteristic *Mtb* lesions. Photomicrographs were obtained from tissue sections representative of the mean values within individual animal treatment groups.

### Statistical analysis

All values are mean ± SD or SEM of individual samples. Data analysis was performed with GraphPad Prism Software (GraphPad Software Inc., version 7.01). The statistical tests utilized have been indicated in respective sections and figure legends.

### Study Approval

The study was approved by the Institutional Biosafety Committee (IBC) and Institutional Animal Care and Use Committee (IACUC) of the (i) Biological Resource Council (BRC), A*STAR, Singapore; (ii) Defence Science Organization (DSO) national laboratories, Singapore, (iii) Colorado State University, USA, and (iv) University of Massachusetts Medical School (UMMS), USA. The study in human healthy volunteers and diabetic patients was approved by the Arnhem-Nijmegen Ethical Committee (NL47793.091.14) and National University of Singapore Institutional Review Board (NUS-IRB #04-140), respectively. Consent from all participants were obtained.

## Acknowledgements

We thank SIgN Flow Facility, ABSL3, BRC laboratory personnel and assistance of Yannick Simoni, Nicholas Ang and Kandhadayar G Srinivasan for assistance. This research was supported by SIgN A*STAR; Immunomonitoring platform grant (#BMRC/IAF/311006, #H16/99/b0/011, and #NRF2017_SISFP09); NIH Grant (#R01HL081149 and #U19AI111224), ERC Advanced Grant (#833247) and a Spinoza grant of the Netherlands Organization for Scientific Research.

## Author contributions

A.S. conceived the idea, designed the study. J.B. performed the experiments and analysed the data, A.L., and M.M. performed the experiments. N.M.G. and H.K. performed mice adoptive transfer and BCG vaccination experiments, and analysed the data. S.L. and E.N. analysed CyTOF data. D.A, J.H.F., A.T. and R.B. performed Guinea pig experiments. J.L. performed RNA-seq studies and B.L. analysed RNA seq data. E.L., T.P.N., A.M.T., A.L., M.G.N., and R.C. provided the clinical samples and related information. A.S. oversaw the study and wrote the manuscript. All authors discussed results and commented on the manuscript.

## Accession Codes

GEO accession of RNAseq data is GSE139948. Primary data is available upon request.

## Supplementary Material

### 1. Supplementary methods

### 2. Supplementary figures

**Supplementary Figure 1.** Phenotyping of Metformin-educated CD8^+^ T cell from naïve wildtype mice in spleen during *Mtb* infection.

**Supplementary Figure 2.** Metformin-treatment expands cluster of memory-like CD8^+^ T cells and CXCR3 expression CD8^+^ T cells in the spleen of wildtype mice.

**Supplementary Figure 3.** *In vitro* treatment with Metformin increases ratio of Ag-specific CD8^+^ TM over TE.

**Supplementary Figure 4.** Metformin-treatment of wild type mice increases CXCR3 expression in the lung CD8^+^ T cells and their subsets.

**Supplementary Figure 5.** Assessment of mitochondrial mass and health, and glucose uptake in CD8^+^ T cells.

**Supplementary Figure 6.** Metformin reprograms host oxidative phosphorylation and immune response in BCG vaccinated mice.

**Supplementary Figure 7.** Effect of metformin treatment on the phenotype of peripheral CD8^+^ T cells from healthy and DM individuals.

**Supplementary Figure 8.** Effect of metformin on long-term *Mtb* clearance and reactivation.

### 3. Supplementary tables

**Supplementary Table 1.** Mouse antibodies used for mass cytometry analysis listing metal conjugate, antibody clone, supplier of each marker

**Supplementary Table 2.** 202 Differentially expressed genes between splenic CD8^+^ T cells from *Cxcr3^−/−^* vs WT untreated mice. FOXO1 targets are depicted.

**Supplementary Table 3.** Enrichment of Canonical pathways (IPA analysis) in 202 DEGs between splenic CD8^+^ T cells from *Cxcr3^−/−^* vs WT untreated mice.

**Supplementary Table 4.** 267 Differentially expressed genes between splenic memory-like CD8^+^ T cells from metformin-treated vs untreated mice.

**Supplementary Table 5.** Enrichment of molecular functions (IPA analysis) in DEGs between splenic memory-like CD8^+^ T cells from metformin-treated vs untreated mice.

**Supplementary Table 6.** 607 Differentially expressed genes between splenic memory-like CD8^+^ T cells from metformin-treated BCG-vaccinated vs untreated BCG-vaccinated mice.

**Supplementary Table 7.** Enrichment of canonical pathway (IPA analysis) in DEGs between splenic memory-like CD8^+^ T cells from metformin-treated BCG-vaccinated vs untreated BCG-vaccinated mice.

**Supplementary Table 8.** Human antibodies used for mass cytometry analysis listing metal conjugate, antibody clone, supplier of each marker.

**Supplementary Table 9.** Socio demographic information of DM cohort and respective different treatment.

**Supplementary Table 10.** Reagents used in this study.

**Supplementary Table 11.** Mouse antibodies and dyes used for flow cytometry listing conjugate, antibody clone, supplier and catalogue number of each marker.

**Supplementary Table 12.** Number of transferred cells into the recipient mice in adoptive transfer experiments.

**Supplementary Table 13.** Human antibodies used for flow cytometry listing conjugate, antibody clone, supplier and catalogue number of each marker.

